# Stochasticity in the Genotype-Phenotype Map: Implications for the Robustness and Persistence of Bet-Hedging

**DOI:** 10.1101/042424

**Authors:** Daniel Nichol, Mark Robertson-Tessi, Peter Jeavons, Alexander R.A. Anderson

## Abstract

For the last few decades modern biology has focused on quantifying, understanding and mapping the genetic characteristics of cells. This genotype–driven perspective has led to significant advances in our understanding and treatment of diseases such as cancer e.g. the discovery of driver mutations and the development of molecularly–targeted therapeutics. However, this perspective has largely ignored the functional outcome of genetic changes: the cellular phenotype. In part, this is simply because phenotypes are neither easy to define or measure as they critically depend on both genotype and context. Heterogeneity at the gene scale has been known for sometime, and there has been significant effort invested in trying to find patterns within it, but much less is understood about how this heterogeneity manifests itself in phenotypic change, i.e. the genotype-phenotype map (GP–map). This mapping is not one-to-one but many-to-many and is fundamentally the junction at which both genes and environment meet to produce phenotypes. Many genotypes produce similar phenotypes, and multiple phenotypes can emerge from a single genotype. To further complicate matters, genetically identical cells in uniform environments still exhibit phenotypic heterogeneity. Therefore a central open question in biology today is how can we connect the abundance of genomic data with cell phenotypic behaviour, this is especially pertinent to the issue of treatment resistance as many therapies act on cellular phenotypes.

Our focus here is to tackle the GP–map question through the use of the simplest functional mapping we can define that also captures phenotypic heterogeneity: a molecular switch. Molecular switches are ubiquitous in biology, observed in many organisms and naturally map molecular components to decisions (i.e. phenotypes). Often stochastic in nature, such switches can be the difference between life or death in environments that fluctuate unpredictably, since they will ensure that at least some offspring are adapted to future environments. For convenience we use Chemical Reaction Networks (CRNs) to define the map of gene products to phenotypes, allowing us to investigate the impact of distinct mappings (CRNs) and perturbations to them. We observe that key biological properties naturally emerge, including both robustness and persistence. Robustness may explain why such bet hedging strategies are common in biology, and not readily destroyed through mutation. Whereas persistence may explain the apparent paradox of bet–hedging – why does phenotypic hedging exist in environments beneficial to only one of the phenotypes, when selection necessarily acts against it? The structure of the molecular switch, itself subject to selection, can slow the loss of hedging to ensure a survival mechanism even against environmental catastrophes which are very rare. Critically, these properties when taken together have profound and significant implications for the emergence of treatment resistance, since the timescale of extinction depends heavily on the underlying GP–map.

## Introduction

Treatment resistance in many diseases is be driven by the pre–existence of resistant phenotypes within the population. Why such phenotypes co-exist (with sensitive phenotypes) and persist in environments never exposed to drug treatment remains a significant unanswered question. Phenotypic heterogeneity has been observed even within isogenic populations of a number of organisms and at many scales [1], from the unicellular – bacteria [2], fungi [3] or cancer cells [4] – through insects [5, 6], plants [7] and even aspects of human development [8]. This inter–cellular variation has been observed even in homogeneous and constant environments, suggesting that aspects of organismal phenotype may be stochastically determined. In environments that fluctuate unpredictably this phenomenon can serve as a survival mechanism by increasing the likelihood that at least some offspring are well–adapted to future environments. Thus, stochastic phenotype determination has been termed *bet–hedging* as a species diversifies the phenotypes within the population in order to “hedge its bets” against future environmental change (see [9] for justification of this naming). Oscillatory environments are common in a range of ecological settings including fluctuating climates, immune–pathogen interactions or cyclic hypoxia within tumours, and the range of phenotypic traits which are thought to display stochastic determination is just as broad.

An important clinical example is that of persister cells which arise stochastically within isogenic populations of infectious bacteria such as *Escherichia coli* [10, 11, 2]. These cells, which constitute a small fraction of the population (< 1%[11]), have reduced metabolism and shut down all non–essential cellular functions. In this dormant state the persister cells are tolerant to the cytotoxic effects of a number of antibiotic agents. Although dormant, these cells can retain the ability to proliferate (although at a drastically reduced rate) and when antibiotic treatment ceases persisters will begin to proliferate, producing non–persisters and driving the re–emergence of the bacterial population. Hence bet–hedging, by creating a small sub–population impervious to those therapies that act on proliferating cells, proves to be an effective survival mechanism against antibiotic treatment. Indeed, bacterial persisters are thought to be a contributing factor to multidrug resistance in a number of diseases [11, 12, 13] and are implicated in the dormancy of chronic diseases, such as Tuberculosis, which can be suppressed but not eradicated [14]. Novel treatment strategies capable of effectively killing persister cells are desperately needed and this need will continue to grow with the increasing incidence of resistance to our presently most effective antibiotics.

Bet–hedging in cancer has been minimally studied; however, a number of aspects of disease course suggest that bet-hedging mechanisms may be important for understanding how tumours evade therapy. In cancer, significant regression of tumours post-therapy leads to a period of remission, followed by the regrowth of aggressive, therapy-resistant lesions. These dynamics can be explained by the classical clonal view of cancer [15], wherein recurring resistant cells are those that happened to acquire the resistance mechanisms through mutation prior to the treatment. However, the high frequency of tumour recurrence in many cancers suggests that therapeutic escape cannot be solely based on mutational luck. Experimental results have shown evidence of transitory resistance [16, 17] indicative of the existence of a small drug–resistant subpopulation that re–establishes a drug–sensitive cancer cell population. Recent experiments have identified the existence such populations of ‘cancer persister cells’ in a cell line of EGFR+ non–small cell lung cancer [18] indicating that bet–hedging may play a role in the emergence of cancer drug resistance [19, 20]. Thus, an understanding of bet-hedging in normal and abnormal (e.g. cancer) cell function may help us understand why certain types of therapies fail while others succeed.

### The Causes of Bet–Hedging

A number of causes of bet–hedging have been identified across different species, but in many cases the precise cause remains an open question. Many bacterial species (see [21] for a review) generate stochastic phenotypic variation by means of *contingency loci* [22] in their genome – short repetitive DNA sequences prone to localised hypermutation and which epigenetically modify the expression of other vital genes. Through hypermutation these loci endow a species with the ability to rapidly generate genetic and phenotypic heterogeneity, forming a quasispecies [23], and implementing a hedging–like strategy.

However, true bet–hedging is characterised by the generation of phenotypic variation without heritable genetic alterations. Beaumont et. al. demonstrated the *de novo* evolution of bet–hedging in the phenotypic trait of colony morphology of the bacterium *Pseudomonas fluorescens* by imposing stochastically fluctuating environments through replating [24]. The molecular mechanism underpinning this switching behaviour was partially elucidated by Gaillie et. al. who identified a single nucleotide change in the gene *carB* as responsible for the emergence of phenotype switching [25]. The precise mechanisms through which this mutation induces phenotype switching remains an open question. Owing to the difficulty in identifying and isolating persister cells, initial attempts to identify similar genetic drivers for bacterial persistence have proved unsuccessful. Following the recent development of persister isolation techniques, a number of contributing genetic factors have been identified (see Lewis [11] for a review). However, whilst over–expression or deletion of these genes were shown to impact the *proportion* of bacterial persisters within a population, none was found to completely inhibit the persister phenotype, suggesting that gene networks and redundancy may be an important part of bet-hedging strategies.

The identification of mutations responsible for the *de novo* emergence of bet–hedging, or for alteration to the proportions of phenotypes in existing hedging populations, offers little insight into the mechanisms responsible for generating the phenotypic heterogeneity. Specifically, what molecular mechanisms allow an isogenic population to produce multiple phenotypically–distinct subpopulations? Current biological thought is that stochasticity or noise in the levels of specific intracellular proteins may drive phenotypic differentiation. Intra–cellular noise can be generated through the stochasticity of gene–expression [26, 27]. For example, variability in gene–expression can be caused by intrinsic random fluctuations in the rates of transcription and translation, by varying abundances of transcription related molecules within the cell, or by the inevitable change in gene copy–number throughout the cell cycle [28]. Further noise is then introduced as molecules undergo Brownian motion within the cell cytoplasm, randomly interacting and introducing asymmetries in their number. These asymmetries can be amplified by feedback motifs within the intra–cellular molecular interaction network. Thus, bet–hedging is thought to arise as the result of noise–driven stochastic switching behaviour in the biochemical processes that underlie the translation of genes to phenotype – the so–called *genotype–phenotype* (GP) map. The nature of these molecular switches, and how they interact with intra–cellular noise to produce consistent phenotypic variation, is poorly understood.

### Mathematical Models of Bet–Hedging

Müller et al. [29] as well as others [30, 31, 32] have used mathematical techniques to demonstrate that bet–hedging constitutes an evolutionary stable strategy (ESS) in stochastically fluctuating environments – offering strictly greater expected fitness than a deterministic one–phenotype strategy under certain constraints on the environmental fluctuations. Further theoretical work by Botero et al. [33] considers when bet–hedging can offer a greater fitness advantage than *phenotypic plasticity*, where phenotypes are modulated via the individual’s ability to sense environmental changes [34]. This previous work derives constraints on the cost of sensing, predictability of environmental fluctuations and the fitness effects of environmental change to determine when bet–hedging, plasticity or determinism is an ESS. These mathematical results are in agreement with the experimental results of Beaumont et al. [24] that demonstrate the *de novo* evolution of bet–hedging, as well as other experiments that empirically demonstrate that bet–hedging offers a fitness advantage in fluctuating environments [35, 36]. There is, however, a disconnect between the mathematical theory of bet–hedging, showing that it is selected for in fluctuating environments, and biological reality, where bet–hedges exist in homogeneous environments. Consider the case of bacterial persisters in *E. coli* in a hospitable environment. Those cells that take on a dormant phenotype reproduce very slowly and hence reduce the average fitness for the population. Thus, there is selection against populations that produce persisters. Despite this selective pressure, persisters remain present in the population and provide a survival mechanism when antibiotics are eventually introduced into the environment. In this paper we propose that this disconnect can be explained by the structure of the GP–map, the complex network of chemical interactions that integrates genetic and environmental factors to produce cellular phenotypes and which lies at the heart of the modern evolutionary synthesis [37].

In previous theoretical work, abstract models of GP–mappings have provided valuable insight into the evolutionary process. Models using computational solutions to predict RNA secondary structure folding have been used to explore the effects of neutral mutations on evolutionary trajectories [38, 39] as well as the evolution of evolvability [40]. Gene regulatory network models have also provided insight into the evolution of evolvability [41]. Gerlee and Anderson [42] used neural networks to study the environmental modulation of phenotypes within a hybrid cellular automaton model of a growing tumour [43, 44]. This model has since found further uses in bridging between the genetic and phenotypic scales [45]. To study the emergence of non–genetic heterogeneity in cancer Huang [46] introduced a conceptual framework in which to understand how complex intracellular regulatory networks shape gene expression profiles and the corresponding cellular phenotypes [47, 48]. In this model gene expression profiles are assigned a “potential” corresponding to the stability of the expression profile to stochastic molecular interactions. This assignment creates an *epigenetic landscape* (in the sense of Waddington [49] – arising not from single genes but the interaction of many). The local minima of potential in this landscape correspond to stable expression profiles (or equivalently in their model, phenotypes) and those gene expression profiles with higher potential will move down–hill through regulatory feedback mechanisms until a stable expression profile is found. Huang argues that non–genetic heterogeneity can then be explained by the existence of multiple accessible stable expression profiles and that phenotypic switching is due to stochastic fluctuations causing jumps between stable states, where the rates of switching depend on the stability or ‘depth’ of the potential wells and external stimuli that can change the magnitude of fluctuations.

Here we develop a model of the GP–map which uses stochastic simulation of simple chemical reactions to mimic the intra–cellular molecular interactions responsible for determining phenotype. Specifically, we stochastically simulate the dynamics of simple multistable reaction networks and assign to each stable configuration a phenotype. These stable configurations are analogous to Huang’s local minima in an epigenetic landscape, but our model differs in that we explicitly build the networks and stochastically simulate their dynamics, as opposed to considering average case deterministic behaviour of a gene–regulatory network. The benefit of this technique is that we are able to directly study the implications of alterations to the network with regard to phenotypic bet–hedging. We demonstrate that the structure of the chemical reaction network directly leads to bet–hedging that is robust to major alterations to the network — offering a possible explanation for the difficulty in identifying single genetic drivers of bet– hedging in many cases. Further, we demonstrate that the network structure can alter the rate of evolutionary convergence to fitness optima and can reduce evolvability, preventing the loss of bet–hedging in homogeneous environments. These findings suggest that natural selection may favour certain chemical reaction network architectures within the GP–map, specifically, those that are robust to alteration or which preserve hedging as a survival mechanism. Finally, we discuss the implications of this result for the evolutionary history of bet–hedging and for the design of treatments for diseases which display non–genetic phenotypic heterogeneity.

## Results

### Chemical Reaction Networks as a Model GP–Map

To investigate the impact of the GP–map on the evolution of bet–hedging we implemented a model mapping with a genetically determined bet–hedge. At the heart of our GP–map is an abstract model of chemical reaction networks (CRNs) that are simulated computationally. CRNs are defined by a collection of labelled chemical species and a list of reactions, with associated rates, between these species. As CRNs explicitly represent biological mechanisms they provide a powerful tool for studying the GP–map with the potential to be closely tied to empirical data. There exist a range of mathematical models of CRNs including continuous models of the mass action kinetics [50], simulation of the stochastic process [51, 52] and analysis using stochastic differential equations [53]. It is entirely infeasible to explicitly model the full array of chemical interactions comprising the translation from genes to phenotypes. We can however investigate smaller CRNs in order to understand what properties the full mapping may possess. This is the approach taken by Cardelli [54] who studied emulation between CRNs – the phenomenon where one network is capable of reproducing the exact mass–action kinetics of another. Cardelli derives algebraic criteria for the existence of emulations from structural properties alone, providing a method to extend studies of simple CRN motifs to larger networks.

A well studied class of chemical reaction networks are those which encode bistable (or multistable) switches [55] in which the series of reactions must eventually terminate in a stable configuration from which no more reactions can occur. The different final configurations of a bistable CRN can be considered different states of a stochastic switch. The probability that the CRN progresses to a specific switch state is dependent on the initial conditions for the network. Here, we assume that cells take one of two phenotypes, A or B, corresponding to the two stable configurations of a bistable CRN (S, R). The phenotype is determined by stochastically simulating the CRN from an initial configuration determined by the genotype according to,

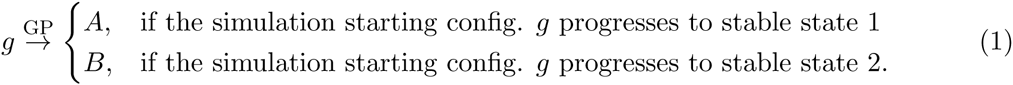

A schematic representation of this model is shown in Figure 1. Depending on the biological system in question these phenotypes could, for example, represent normal/persister cells in *E. coli*, the Cap^+^/Cap^-^ cells in colonies of *P. fluorescens* or the lysis/lysogeny decision switch in bacteriophage lambda. We assume that convergence time to an equilibrium state in the CRN is sufficiently fast (in comparison to the cell cycle timescale) that we may take it to be instantaneous.

**Figure 1:**
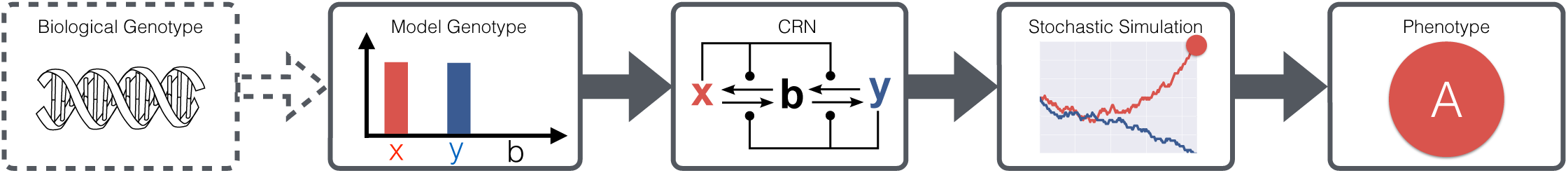
Schematic representation of the chemical reaction network model for the genotype–phenotype mapping. To reduce the complexity of our model the genotypes are abstracted away from biological reality (dashed lines).

We define the genotypes, *g*, in our model to be the initial numbers of the chemical species within the chemical reaction network. This definition of genotype is an abstraction in which we choose to ignore the physical mechanisms of inheritance and expression and instead consider genotypes from the perspective of the expressed gene products. We choose not to model the inherent stochasticity in gene expression, a phenomenon previously associated with bet–hedging, in order to explore the implications of the stochasticity of interactions induced by Brownian motion of molecules within the cell cytoplasm. For simplicity we model genotypes as the initial and instantaneous expression of two gene products *x* and *y*, denoted by *x*_0_ and *y*_0_ respectively, and we assume all other species within a CRN are initially 0. Mutations are modelled as changes to the initial abundance of *x*_0_ and *y*_0_ and we assume that there exists a maximum abundance that *x*_0_ or *y*_0_ can attain. In particular, for an individual with genotype *g* = (*x*_0_,*y*_0_) the possible mutants are given by

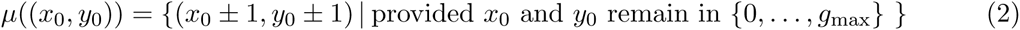

We assume that mutations occur at replication with some probability *μ* and that where more than one mutation is possible each is equally likely. Finally, for our long-term evolutionary simulations (the section entitled “Evolutionary Loss of Bet-Hedging”) we will further impose the simplifying restriction that *x*_0_ + *y*_0_ = *g*_max_.

There are numerous examples of bistable switches that can serve as genotype-phenotype mappings within this framework. Examples include the Approximate Majority, Direct Competition and Great Wall Kinase switches studied by Cardelli and Csikász-Nagy [55] and Cardelli [54]. Here, we utilize a number simple bistable CRNs to serve as examples demonstrating the impact of the GP-map on the evolution of bet-hedging. Each of our model switches is derived from the CRN shown in Figure 2 by modification to the rates of the individual reactions. Figure 3 shows the relationship between the model genotype and the probability of stochastic determination to phenotype *A* for four such molecular switches - the Direct Competition (DC) switch, a version of this switch biased by a preference to produce the species *x* (DCx), a version biased to produce the species *y* (DCy) and the Approximate Majority (AM) switch studied by Angluin et al. [56] and later Cardelli and Csikász-Nagy [55]. By picking appropriate initial conditions (i.e., the values of the gene products inherited by cells) for the chemical reaction networks, we can approximate any switching probability and equivalently any proportion of either phenotype arbitrarily closely (Figure 3 Row 3). It follows that bet-hedging is an extremely simple biological mechanism, one that can be produced by the stochastic interactions between as few as two or three molecules. This simplicity could help explain the ubiquitous nature of stochastic phenotype determination and why a large number of genetic factors have been implicated in this process across species; as simple molecular switches suffice they are more likely to emerge through mutation and are an effective long-term survival strategy.

**Figure 2:**
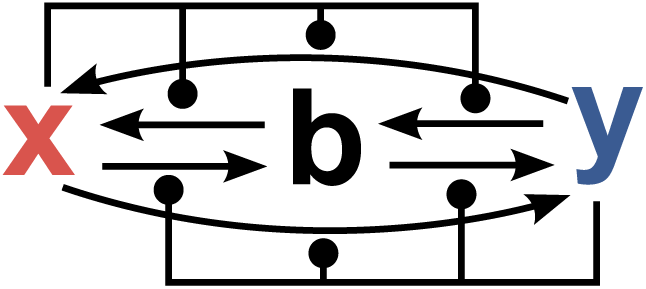
The CRN from which our example switches are derived. Our example switches are created by modifying the rates of each of the six reaction that build this CRN.

**Figure 3:**
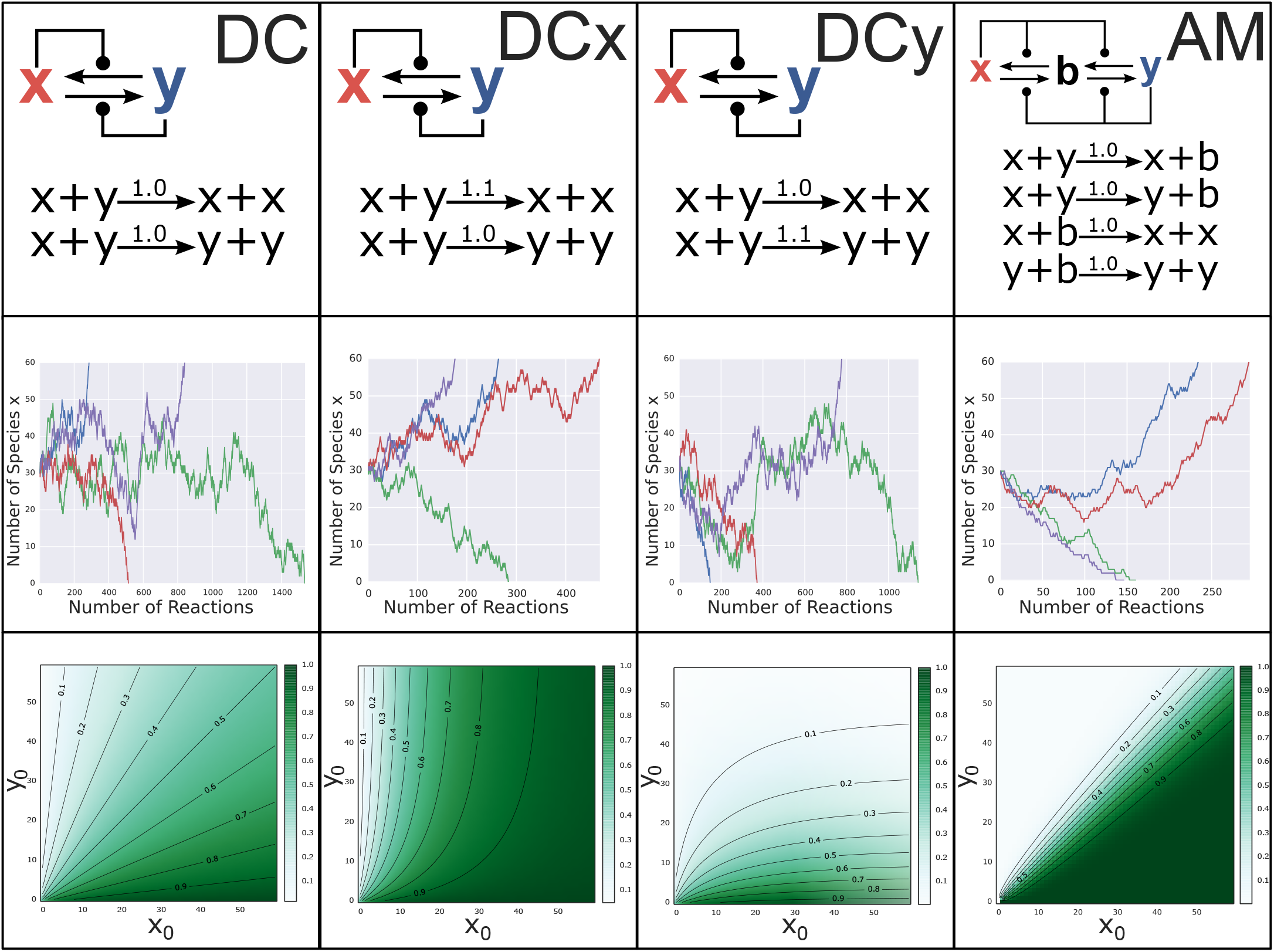
Example molecular switches as GP–Maps. Each column shows the characteristics of one of the four switches (Direct Competition, *x*–Biased Direct Competition, *y*–Biased Direct Competition and Approximate Majority) introduced in the main text. The first row shows the name, chemical reaction network structure and precise definition of each switch. The second row shows stochastic trajectories of the number of molecule *x* in the system for four different simulations of each switch. The starting condition in all simulations is *x* = *y* = 30, (*b* = 0 for the AM network). Note that all of the switches are able to resolve to either of the stable conditions, *x* = 60 or *x* = 0, which correspond to the phenotypes *A* and *B* respectively. Row three shows contour plots displaying the probability of switching to phenotype *A* for each possible initial condition with 0 < *x*_0_, *y*_0_ ≤ 60 and *x*_0_ + *y*_0_ ≠ 0 (*b* = 0 for the AM switch). Contour lines show subspaces of genotype space of equal hedging probability for hedges equal to 0.1, 0.2,…, 0.9.

### Robustness and Redundancy in Molecular Switches

The CRN can demonstrate how redundancy in the network can implement robust molecular switches that maintain bet-hedging even when individual components are removed. Figure 4 shows a version of the simple DC switch from Figure 3 in which the species *x* and *y* are duplicated. In this network, which we call DCdup, the set of stable configurations are determined by *x*+*x′* = 0 or *y*+*y′* = 0. If we associate the phenotypes *A* and *B* with these two configurations respectively then the switching probability on initial conditions (*x*_0_+*x*′_0_, *y*_0_, *y′*_0_) is identical to the switching probability of DC with initial conditions (*x*_0_ + *x′*_0_, *y*_0_ + *y′*_0_) (a simple mathematical argument to establish this proceeds by symmetry and re-labelling the species). The advantage of DCdup is that it maintains its switching properties even if chemical species are removed. Figure 4 shows numerical solutions for the CRN switching probability when the species *x* is deleted (middle network) and then when both *x* and *y* are deleted (right hand network). These induced CRNs maintain switching behaviour similar to the original CRN DCdup. The network induced by deleting *x* (or by symmetry *y*) behaves precisely as DCdup with initial condition (*x*_0_,0,*y*_0_, *y′*_0_) (by symmetry (*x*_0_, *x′*_0_, 0,*y′*_0_)). Further, removing both *x* and *y* from DCdup creates a version of the DC switch in the species *x′* and *y′* which behaves precisely as the DCdup switch on initial conditions (0, *x′*_0_,0, *y′*_0_). This means that deletion of a single chemical species will change the switching if the numbers of all other species (in our evolutionary model, the genotype) are held constant. An example is shown by the red circles in Figure 4 where deletion of chemical species shifts the switching probability whilst maintaining a hedge. The resulting switch has the potential, after adjustment to the initial conditions, to precisely replicate the switching behaviour of the larger network. It follows that the DCdup is robust to the removal of chemical species - a mutational event that in our model can be interpreted as a gene deletion or mutation that down-regulates the expression of a gene. Reversing our previous argument, the switching behaviour of the CRNs in Figure 4 show how the DC switch is robust to gene duplications or upregulating mutations. The duplication of *x* or *y* (or both) produces a CRN which is still a molecular switch, although with a possibly altered switching probability depending on the initial numbers of the species.

**Figure 4:**
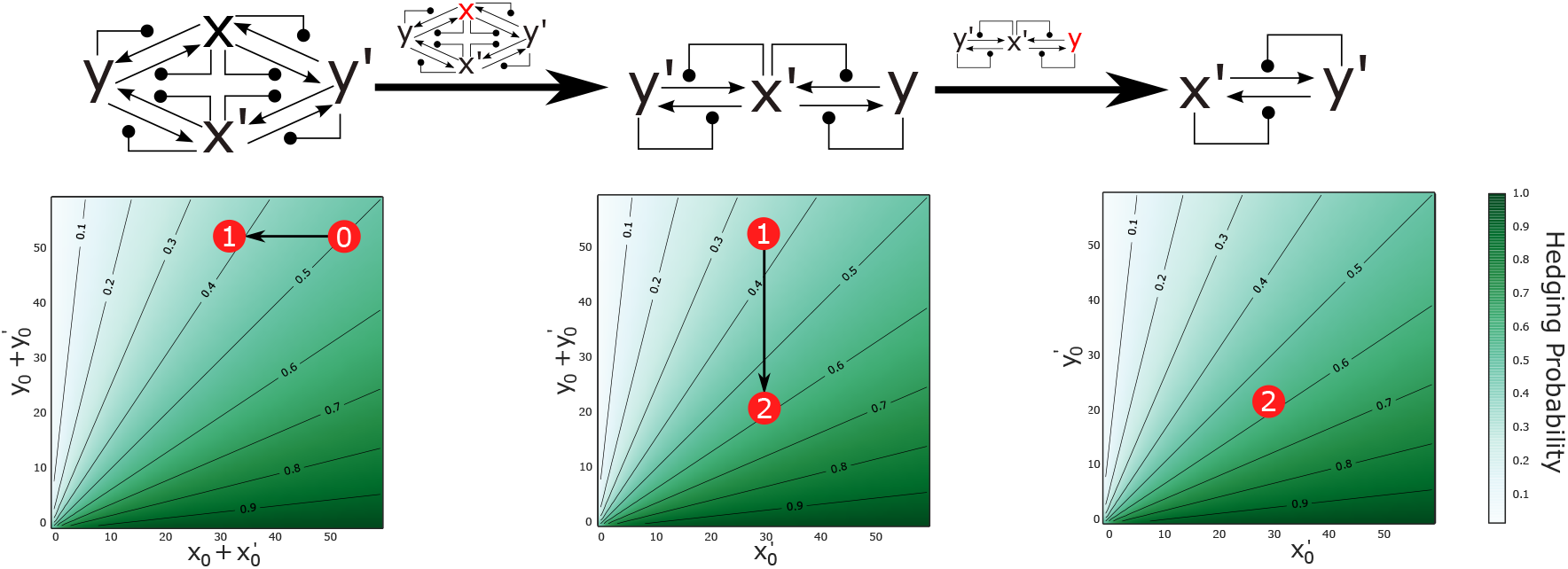
Redundancy in the chemical reaction network implementing the DC switch maintains molecular switching when chemical species are deleted. Marked in red is the switching probability for initial conditions (20, 30, 30, 20) before deletion (0), after the deletion of *x* (1) and after the deletion of *x* and *y* (2). Note that these deletions maintain the stochastic switch but alter the switching probability. Contour lines show initial conditions of equal switching behaviour.

A similar implementation of the AM molecular switch is shown in Figure 5 and is robust to the removal of species, *x, y* and *b* in any order. As with the DCdup network, the removal of any chemical species will change the switching probability, shifting the proportion of each phenotype in the population, but will not entirely prevent switching. Finally, we note that molecular switches can also be robust to the alteration of reaction rates, including the entire removal of reactions. Specifically, we note that each of the example switches presented in Figure 3 are modifications of the larger CRN shown in Figure 2. The observation that we can build molecular switches that are robust to the deletion or duplication of species, and to the alteration of reaction rates, provides insight into the long standing failure to identify single genetic factors responsible for bet-hedging. It may be that bet-hedging arises not from a single factor but from the interactions of many, none of which are individually necessary to maintain the switching behaviour.

**Figure 5:**
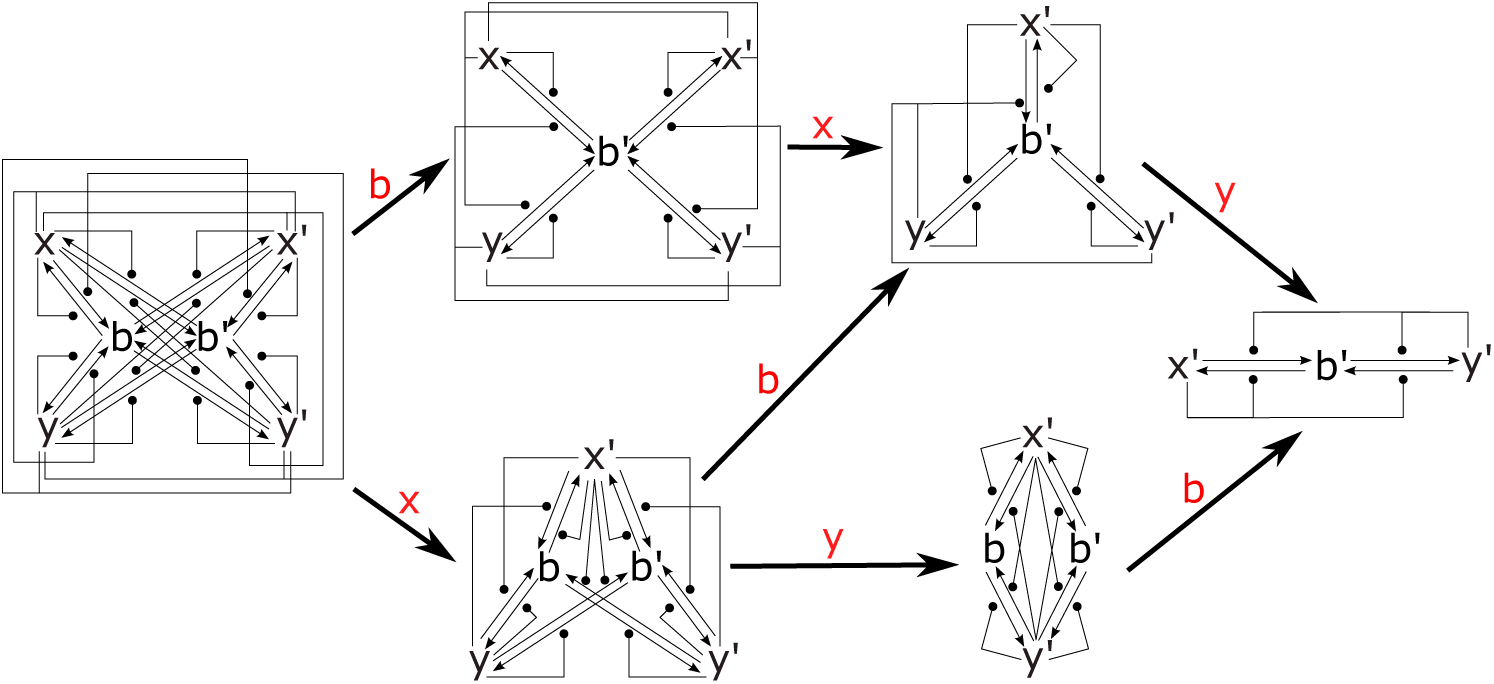
Redundancy in the chemical reaction network implementing the AM molecular switch. Switching is maintained if the species *x, y* and *b* are removed in any order. We omit the case where *y* is removed before *x* due to symmetry.

### Evolutionary Loss of Bet–Hedging

Each molecular switch we study has the ability to (approximately) produce any arbitrary switching probability between phenotype *A* and *B*. Thus, for a given population genotype, the specific switching mechanism responsible for stochastic phenotype determination is irrelevant. However, over longer timescales the structure of the CRN responsible for switching has a significant effect on the evolution of bet-hedging. In our model the probability that a genetic mutation (modelled as a change to the initial conditions of the CRN) fixes within an existing population is determined by the new switching probability, and the associated expected fitness, of that new mutant genotype. Here we consider the rate at which evolutionary loss of the bet-hedging occurs in a fixed environment strongly favouring one phenotype (*A*) over a second (*B*) to demonstrate the importance of the GP-mapping on predicting the evolutionary fate of bet-hedging.

We simulate evolution by assuming a large asexually reproducing bet-hedging population exists under Strong Selection Weak Mutation (SSWM) dynamics. The population is assumed to be isogenic and the population genotype is periodically replaced by a fitter mutant. For simplicity, and to avoid issues regarding the phenotypic/fitness implications of switching on fewer molecules, we enforce that *x*_0_ + *y*_0_ remains constant (*x*_0_ + *y*_0_ = *g*_max_ = 60 for computational efficiency) throughout our simulations. Thus, mutations consist of simultaneously incrementing *x* (or *y*) and decrementing *y* (resp. *x*). The genotype is then entirely determined by *x*_0_ ∈ {0,…,60}. Our simulation proceeds by repeatedly generating a mutant of the current population genotype and computing the probability (using the theory of multi-type branching processes, see the Materials and Methods) that this mutant fixes as the new population genotype. Throughout the remainder of this work we take the two phenotypes *A* and *B* in our model to correspond to a high fitness, proliferative phenotype and a low fitness, slow proliferating phenotype respectively, to mirror the phenomenon of bacterial persistence. We parameterise our model by assigning to phenotype *A* (respectively *B*) a fitness value *w*_*A*_ (resp. *w*_*B*_) encoding the expected number of offspring an individual of that phenotype will have over a single timestep in our population dynamics model (see Materials and Methods).

The invasion probability is computed independently of the (assumed to be large) population size and is dependent only on the (stable) distribution of phenotypes in the population at equilibrium. For this reason we need only know the relative fitness values of the phenotypes. Hence we may take the timesteps of population dynamics to be equal to the expected division time of a cell of phenotype *A*. Mirroring the scenario of persistence in *E. coli* we take this timestep to be *t* = 60mins and *w*_*A*_= 2.0. The reproductive rate of persister-type cells is unknown and thus, in order to match their behaviour qualitatively, we take our persister-like phenotype cells to reproduce at a rate 10 times slower than the proliferative phenotype and set *w*_*B*_= 1.01. Although this parameterisation is only an approximation to the genuine population dynamics of persister cells it is sufficient as an illustrative model demonstrating the importance of the genotype-phenotype mapping on the evolutionary dynamics of bet-hedging.

Figure 6 shows how changes in the population genotype manifest themselves as changes in the average population fitness. The expected population fitness increase associated with the mutation of the current genotype *x*_0_ to *x*_0_ +1 is not equal for all x_0_ and is dependent on the GP-map. Thus, the invasion probabilities for mutations differ depending on the current population genotype and the form of the GP-map. Figure 7 shows the probability of a single mutant genotype *x′*_0_ invading a resident population of genotype *x*_0_ in a hospitable environment. This probability is dependent on the associated increase in average population fitness, as shown in Figure 6. Note that, in the hospitable environment, only mutations which increase the proportion of phenotype *A* are beneficial and hence, as our invasion probabilities are determined from the theory of branching processes, are the only mutations that can fix. Figure 8 shows the probability of successive beneficial mutations, *x*_0_ + 1, invading a resident population of genotype *x*_0_. In this figure we see the impact of the genotype-phenotype map on the evolutionary dynamics. For the DC, DCx and AM switches the probability of the next beneficial mutation fixing reduces for each successive mutation. The magnitude of this decrease is dependent on the switch and in the case of DCx and AM approaches 0. Conversely, for the DCy switch each successive mutation is more likely to fix.

**Figure 6:**
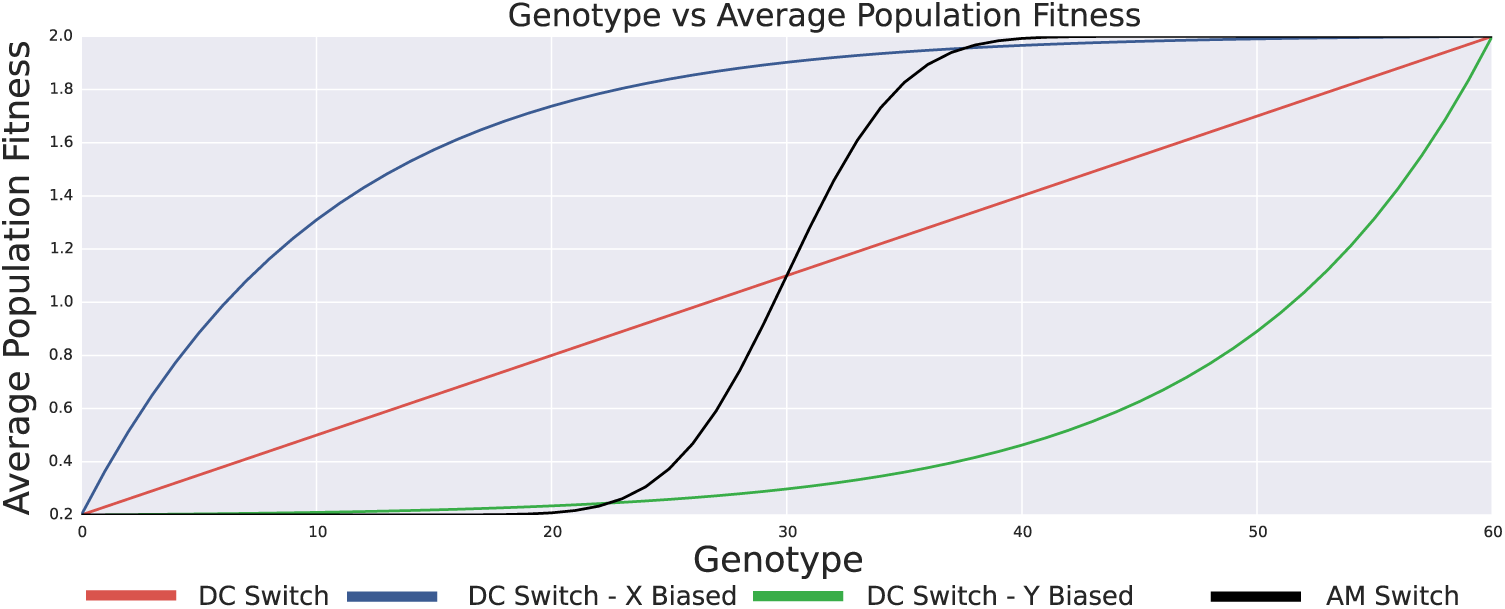
Relationships between the genotype and the associated average population fitness.

**Figure 7:**
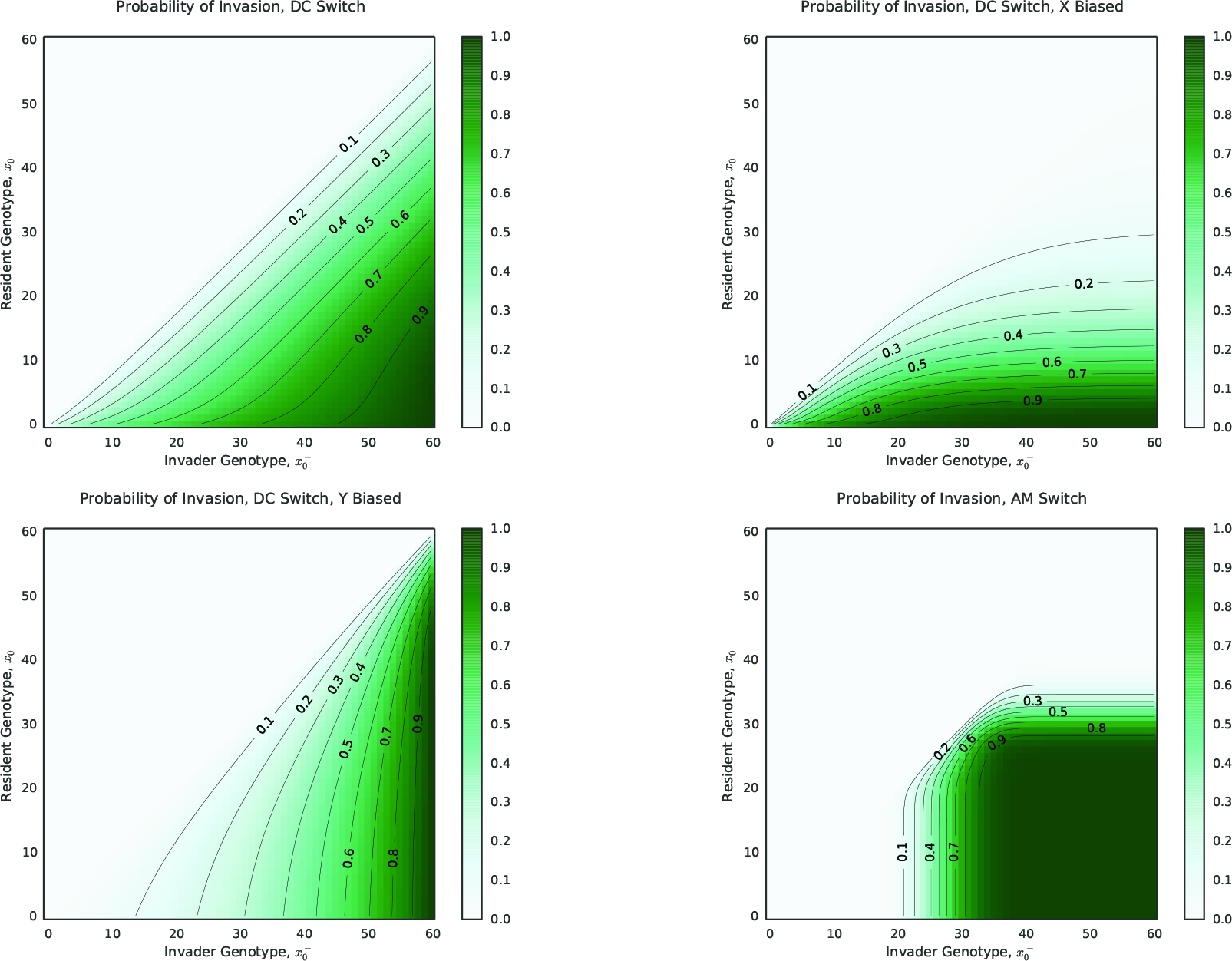
Invasion probabilities for resident and invader genotypes. The invasion probability of an individual of genotype *x′*_0_ into a resident population of genotype *x*_0_ for each of the molecular switches. The probabilities are computed using the theory of branching processes (see Materials and Methods) for fitness values *w*_*A*_ = 2.0, *w*_*B*_ = 1.01.

**Figure 8:**
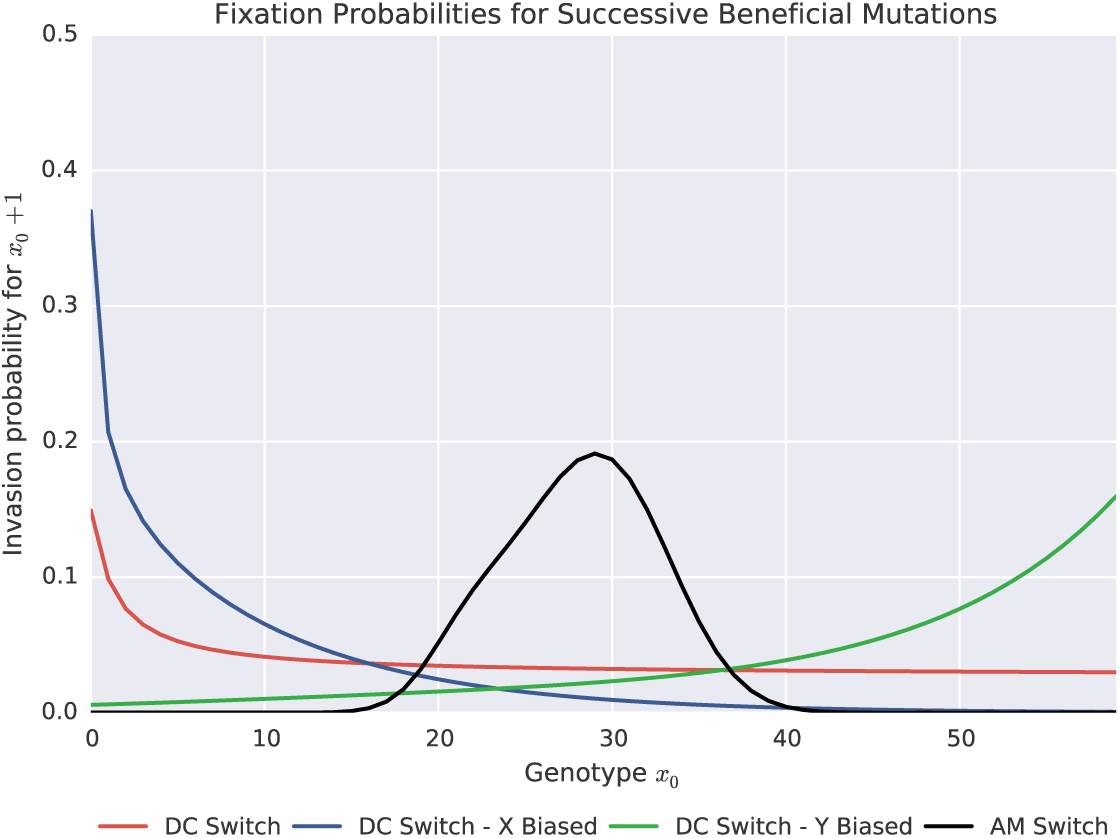
The fixation probability of subsequent mutations differs for each GP–map. For each of the three CRN networks we study the invasion probability of a beneficial mutant *x*_0_ + 1 into a population with genotype *x*_0_ is highly dependent on the GP–map. For the DC switch this mutation always offers a significant fitness advantage and has non–zero invasion probability. For the AM switch this mutation offers a large fitness increase for intermediate values of *x*_0_ but it essentially neutral for extremal values. As we assume a large population the invasion probability is thus approximately 0.

We next consider the evolutionary trajectories of bet-hedging populations endowed with each of our sample GP-maps, DC, DCx, DCy and AM from an initial genotype corresponding closely to a 50% hedging probability, in the hospitable environment which strongly favours phenotype *A*. As the DC and AM switches are symmetric the genotype corresponding to a 0.5 hedging probability is *x*_0_= 30. For DCx the closest genotype to a 50% hedge is *x*_0_ = 7 which corresponds to a probability of 0.49. For DCy the closest genotype to a 0.5 hedging probability is *x*_0_ = 53 and corresponds to a probability of 0.51. As deleterious and neutral mutations cannot fix under our model of population dynamics, the population genotype will be periodically incremented until x_0_ = 60 and the bet–hedge is lost. Figure 9 shows the evolutionary trajectories towards the loss of bet–hedging for populations endowed with each of the candidate GP–maps (Figure 3). The convergence dynamics for these populations is very different. For the DC, DCx and DCy, switches the expected convergence times can be determined as the expectation of a sum of non–identical independent geometric distributions and is given by the sum of the reciprocals of the fixation probabilities of the successive mutations. We find that the expected number of mutational events required for a complete loss of bet–hedging are given by

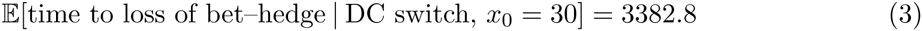

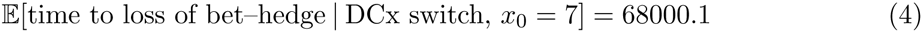

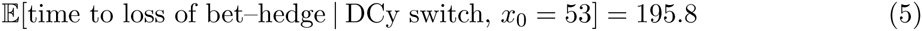

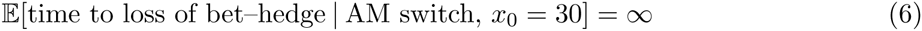

**Figure 9:**
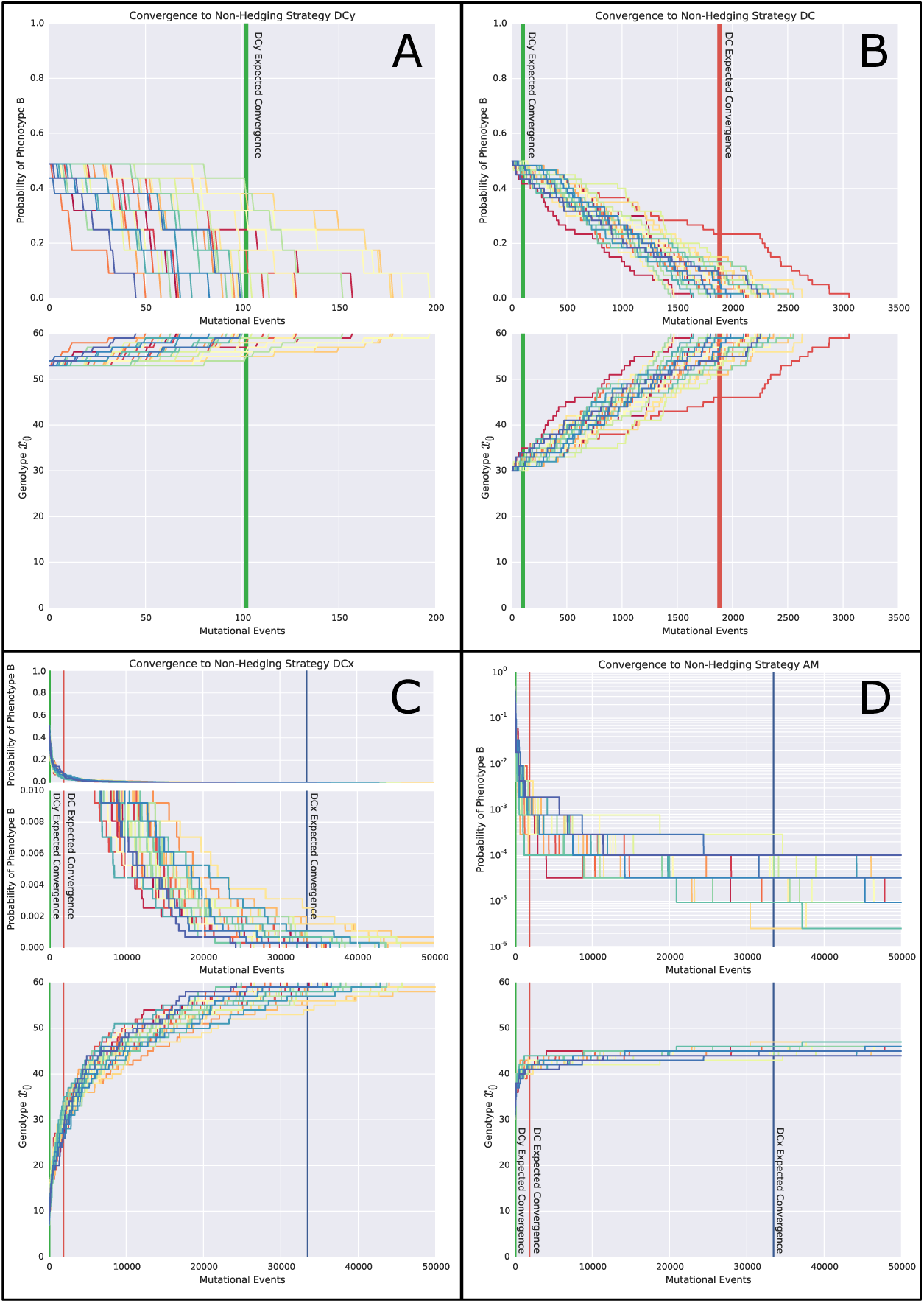
Convergence dynamics through genotype and probability space for the GP– maps defined by DC, DCx, DCy and AM. 30 stochastic realisations of the evolutionary simulation through both genotype and probability space are shown for A) The DCy switch. B) The DC switch. C) The DCx switch. D) The AM switch. Due to the rapid initial change in hedging probability for the DCx switch, the convergence dynamics are also shown on a restricted scale. As the probability of phenotype B rapidly approaches 0 in the AM switch simulation but never converges, the dynamics are shown on a logarithmic scale. The expected convergence time for the DCy switch is marked in green, for the DC switch in marked in red and for the DCx switch is marked in blue.

In the case of the AM network, each subsequent mutation provides a diminishing increase in fitness until mutations are approximately neutral. The probability of neutral mutations fixing within our model of invasion dynamics, which models the population size as tending to infinity, is 0. In reality, the actual convergence times in the AM will depend on the population size. For large populations, as is our assumption, the timescales will be sufficiently long that we take it as equivalent to the evolutionary trajectory never converging ^1^.

### Simulation of Therapeutic Intervention

To demonstrate the importance of the underlying molecular switch in determining an effective treatment strategy for diseases with bet–hedging driven resistance, we implemented a non– spatial, individual–based model simulating the effects of drug treatment on a bet–hedging population. As above, we assume the existence of two phenotypes: one proliferative, but drug sensitive, phenotype *A*; and one drug–resistant, slowly–proliferating, phenotype *B*. Our model takes the form of a discrete time–step death–birth process in which population dynamics are simulated in either of two environments – a hospitable environment (as in our long–term simulation of evolutionary loss of bet–hedging) and a drug–treated environment. The details are provided in the Materials & Methods. We associate both a death rate and a birth rate with each phenotype in order to simulate stochastic extinction dynamics under drug treatment and assume a single discrete timestep corresponds to 1 hour. In the hospitable environment we assign a reproductive rate 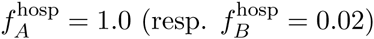 to individuals of phenotype *A* (resp. *B*) and assume the death rate 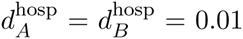 is equal for both types. These parameters ensure that the overall fitness of each phenotype, 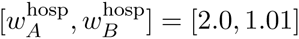, matches the fitness values used above. Finally, for the drug–treated environment we assume the parameters remain unchanged except for an increased death rate for individuals of phenotype 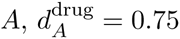.

For each of the molecular switches presented in Figure 3 we performed a simulation of therapy following a treatment holiday. For different timescales, measured in the number of mutational events varying from 0 to 100000, we used our evolutionary simulation to determine the expected population genotype after a holiday from treatment over that timescale. In each case the initial genotype was chosen to correspond as closely as possible to a 50% hedge. We then simulated continuous drug treatment on a population of 10^10^ bet–hedging cells with the hedging probability determined by this pre–calculated expected population genotype. For each treatment holiday length and each molecular switch we performed 2000 simulations of treatment and recorded the time until extinction. Mutations were not permitted during these simulations. The extinction times are presented as histograms in Figure 10. The timescale of holiday required to reduce the extinction time to correspond to a viable treatment length (marked in the figure by a star) varies by orders of magnitude dependent on the form of the molecular switch and associated GP–map.

**Figure 10:**
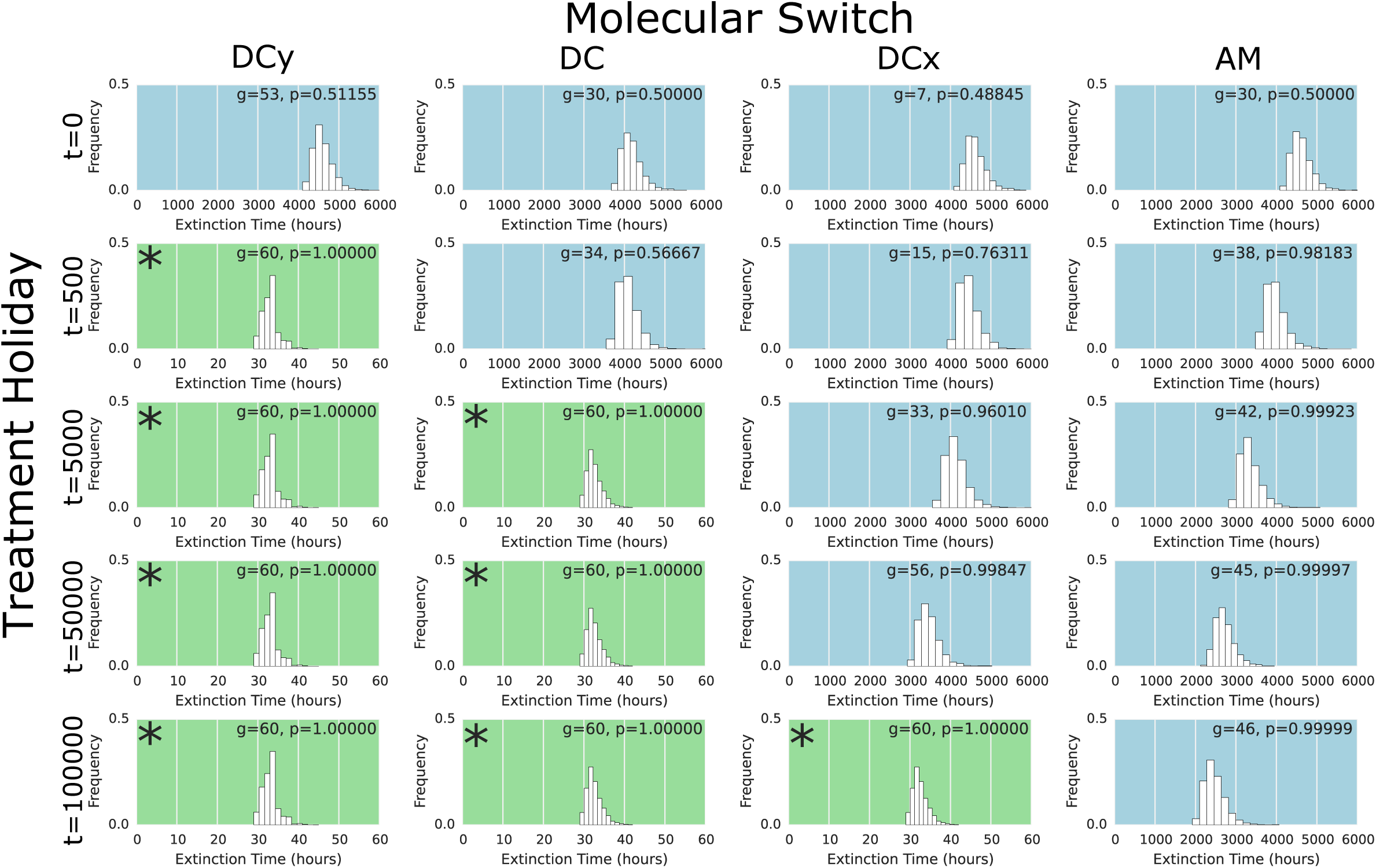
Extinction dynamics for populations endowed with different molecular switches over different timescales of treatment holidays. Each histogram shows the distribution of extinction times over 2000 simulations of treatment in an individual–based model. The molecular switch used as the GP–map is shown as the column heading. The genotype and associated probability of phenotype A (shown inset to each subfigure) are determined by an evolutionary simulation of a treatment holiday for a timescale determined by the row. A blue background indicates extinction times longer than a timeframe viable for an antibiotic treatment, a green background (or inset star) indicates extinction times within this timeframe.

These simulations demonstrate the importance of understanding the underlying molecular mechanisms that drive phenotypic bet–hedging. Specifically, we see that the length of treatment required to drive a population to extinction is dependent on the proportion of each phenotype in the population, which is in turn determined by the population genotype and molecular switch. The length of treatment holiday required to steer the evolving population into a more rapidly treatable configuration, a sufficiently small proportion of persister–like individuals to permit rapid extinction, is dependent on the underlying molecular mechanism. Thus, to understand and predict the efficacy of treatment holidays as a potential therapeutic intervention for a disease with bet–hedging driven resistance it is imperative that we understand the underlying GP–mapping.

## Discussion

We have introduced a model for the genotype–phenotype map which uses stochastic simulation of chemical reaction networks to determine phenotypes. Whilst other models of the GP–map have modelled the emergence of non–genetic phenotypic heterogeneity using network models, namely those of Gerlee and Anderson [42] and Huang [46, 48], ours is the first to explicitly model the stochastic aspect of phenotypic determination at the level of the molecular mechanism. Using this model we have demonstrated how remarkably simple chemical reaction networks can implement complex switching behaviour and produce populations in which different proportions of cells, determined by the initial conditions of the network, take on different phenotypes. Our model, whilst abstract in its representation of chemical reactions, is closely related to the mechanisms of gene translation and the subsequent reactions that govern intracellular regulatory networks and, as such, has the potential to provide valuable insight into the mechanisms responsible for the stochastic determination of phenotypes. Further, through this mechanistic understanding we are able to gain insight to how the phenomenon of bet–hedging evolves under Darwinian selective pressures.

We have demonstrated how introducing redundancy, a common feature of many biological systems, into the chemical reaction networks governing the GP–map can create molecular switches that are robust to the addition or removal of chemical species. Assuming that the chemical species of our CRN correspond to expressed gene products, this finding shows how redundancy, which can arise through neutral or nearly–neutral mutation events, can ensure that the phenomenon of bet–hedging is not lost through gene deletions or duplications. Critically, this observation may explain the failure to identify genes responsible for bacterial persistence. For example, in the review by Lewis [11] mutations to the genes hipA, rmf, sulA, and toxi-nantitoxin (TA) loci relBE, dinJ and mazEF were highlighted as possible drivers of bacterial persistence. However, deletion of rmf, relBE or mazEF has been demonstrated to have no effect on the phenomenon of persistence, owing possibly to redundancy in TA modules, whilst deletion or over expression of hipA can change the proportion of bacterial persisters but not eradicate them. This is consistent with our predictions, since deleting any single species in the CRN will not prevent bet–hedging or non–genetic heterogeneity, instead it will only alter the switching probability, or at the population scale, the proportions of each phenotype. In the context of bacterial persistence, the conclusion taken from the experiments reviewed in Lewis [11] need not be that none of the potential genetic factors identified are the ones driving bacterial persisters, instead it may be that the search for a single genetic factor responsible for bet–hedging is doomed to fail. Indeed, it may be that bet–hedging emerges from the interactions of a collection of genetic factors in the sense of the epigenetic landscape introduced by Brock et al. [47] and Huang [46]. If this is the case, then to identify the biological mechanisms responsible for hedging we will need to move beyond the gene–centric perspective and begin to identify those interactions responsible for stochastic determination of phenotypes.

Mutations in cancer have often been associated with their direct effect on phenotypes; the concept of a driver mutation is that it creates a new phenotype that is distinctly more fit than the non-mutated variant, leading to clonal expansion [57]. However, the results presented here suggest another phenomenon, in that mutation of one or more genes that feed a CRN– based bet–hedging mechanism need not induce novel phenotypes, but simply alter frequencies of pre–existing phenotypes within the population. This change in phenotypic ratio could have implications for cancer progression, even in the absence of any novel phenotypes. Consider the phenomenon of tumourigenic cells, where it is thought that only tumour cells of a certain phenotype can form a growing mass [58, 59, 60]. Clonal genetic heterogeneity can explain the existence of a tumourigenic subpopulation if certain driver mutations were responsible for the tumourigenic phenotype. However, an alternative mechanism is that stem or stem-like tumour cells in the population give rise to a population of heterogeneous phenotypes. In the traditional stem-cell model, a phenotypic hierarchy exists where the stem cell produces the range of tumour cell phenotypes [61]. Under this model, cancer stem cells divide to produce either more cancer stem cells (self–renewal) or cells with non–stem phenotypes down the hierarchy. This cellular decision is often taken to be stochastic (an example of bet–hedging) and thus our results highlight the potential for mutations to alter the probabilities of self–renewal or differentiation, which have been shown to have signficant impact on many aspects of tumour progression [62].

An alternative bet–hedging mechanism for tumourigenicity is that the tumourigenic phenotype is transient and stochastically determined (and potentially influenced by the microenvironment). Evidence for this phenomenon is highlighted in recent work by Quintana et al. [63] that shows that the tumourigenic potential of individual melanoma cells is similar despite high heterogeneity of many markers on the initialising cell. No driver or stem population was found, and indeed, the heterogeneity of marker expression was recapitulated by most tumourigenic cells, regardless of the starting pattern of expression. Such a mechanism would have different implications from those presented above, because targetable cells would be much more difficult to define, but the predictions of our model remain the same: that genetic mutations can shift the frequency of tumourigenic phenotypes and profoundly impact cancer progression.

We have further demonstrated that the structure of the chemical reaction network governing the GP–map has important implications for the evolutionary loss or gain of bet–hedging. By considering mutations that shift the mean expression levels of genes, we find that the time taken for a 2–phenotype bet–hedge to be lost in an environment favourable to only one of the phenotypes can vary by orders of magnitude depending on the GP–map. Thus, if bet–hedging is a survival mechanism in the event of rare catastrophic environmental change (e.g., bacterial persistence in the event of antibiotic exposure or treatment resistance in cancer as a result of targeted therapy) then the GP–mapping can prevent loss of this survival mechanism over the long timescales in which catastrophe does not occur. Specifically, the mapping from an abstracted genotype (modelling the gene expression profile) to the probability of an individual taking each of the two phenotypes is determined by the structure of the molecular switch underlying the GP–mapping and can change the rate at which evolution converges to a fitness optimum. If a bet-hedge is implemented by a switch such as the Approximate Majority (AM) network (Figure 3) then each successive mutation towards a one phenotype strategy induces a diminishing increase in the probability of generating that phenotype. Thus, in a constant environment favouring one phenotype over another, each subsequent mutation induces a diminishing increase in average fitness until it is essentially neutral. As neutral mutations fix with likelihood inversely proportional to population size, the structure of this molecular switch substantially slows convergence to a one–phenotype strategy. Alternatively, if a different switch determines the GP–map, for example the DCy switch presented here, the convergence can be orders of magnitude faster as the fitness benefit of each subsequent mutation increases. This result provides a possible solution to the apparent paradox of bet–hedging – why does hedging exist in environments beneficial to only one of the phenotypes when selection acts against it? The structure of the molecular switch, itself subject to selection, can slow the loss of hedging to ensure a survival mechanism even against environmental catastrophes that are very rare.

These results have important implications for previously suggested theoretical treatment strategies for diseases that display bet–hedging–driven drug resistance. Theoretical strategies suggested to combat bet–hedging–induced multi–drug resistance focus on identifying novel agents capable of killing persister cells and identifying genetic (or downstream) mechanisms that can be targeted to prevent the persister phenotype from emerging. This latter strategy bears a striking resemblance to the targeted therapy revolution in the treatment of many cancers. The identification of molecular targets which when inhibited induce cell death led to the discovery of a number of targeted therapies for melanoma, non–small cell lung cancer and colorectal cancers. These drugs are, in the short term, remarkably effective, however the effects are rarely durable. Mutations that abrogate the effects of targeted therapies quickly emerge during treatment, driving resistance and ultimately mortality. To improve the efficacy of targeted therapies (as well as traditional chemotherapeutics) we must consider how selective pressures induced by therapy drive Darwinian adaptation. The results of our chemical reaction model shed light on this Darwinian adaptation and suggest that targeted therapies to prevent bet–hedging may either be impossible, or where they do exist, prone to fail due to the re–emergence of bet–hedging through evolution. More precisely, the discovery of a single “silver bullet” genetic factor which when targeted can switch off multi–drug resistant dormant phenotypes is unlikely, owing to redundancy in the network architecture. As bet–hedging offers an effective survival mechanism, a fact supported by its ubiquitous role in survival across many species, and as the mechanisms responsible are predicted by our modelling to be remarkably simple, it should be unsurprising that robustness has evolved to maintain it. Of course, this does not rule out the potential of targeted therapies entirely. It may be possible to identify multiple targets which when simultaneously targeted prevent hedging. Alternatively, targets may be identified that shift the proportion of resistant or dormant individuals within a population to a manageable level, either permitting treatment with other cytotoxic agents or allowing us to drive disease into a dormant or even extinct state.

A second theoretical treatment strategy suggested in the design of multi–drug therapies for cancers, as well as those for highly resistant infections [64], is the introduction of treatment holidays. The traditional doctrine for therapy is that we should treat diseases using the most potent drug with the highest tolerable dose until the disease is cured. Mathematical evolutionary models of disease progression suggest that this approach may actually drive the emergence of resistance. Here we implemented an individual–based model of the dynamics of a bet–hedging population under treatment to explore the efficacy of treatment holidays. Coupled with a long–term evolutionary simulation to model treatment holidays we explored the impact of the molecular mechanism driving bet–hedging on the efficacy of treatment holidays. Our model suggests that it is the GP–map, and in particular how it hinders or promotes the rate of evolutionary convergence, that holds the answer to whether treatment breaks can undo the emergence of drug resistance. If evolutionary loss of bet–hedging is fast, as it would be if the molecular mechanism responsible resembled the DC or DCy explored in this paper, then short to medium term treatment holidays could be expected to steer evolution such that bet–hedging is lost from the population. However, if instead the underlying mechanism resembles the AM switch then we would know that a treatment break is unlikely to undo multidrug resistance induced by bet–hedging within a timeframe relevant to disease progression. Finally, we note that interfering with the underlying switching mechanism, for example through the use of targeted therapies, could alter the switch properties sufficiently that the targeted therapy administered alone (in the absence of cytotoxic agents) could drive evolution to remove hedging from the population. An example of such a switch is presented in Figure 2 where removal of the species b induces a switch equivalent to the DC switch, which is susceptible to a treatment holiday in a short period of time.

In this paper we take the initial conditions of our chemical reaction networks to be genetically determined, allowing us to explore the implications of the structure of the GP–map on the evolution of bet–hedging. The chemical reaction model could also be used to study additional aspects of the evolution and effects of non–genetic heterogeneity other than those presented here. For example, we could instead have taken the initial conditions in the CRN to be environmentally determined, through the concentration of some diffusible factors in a spatial domain, and the network itself to be genetically determined. In this case the CRN model would give a GP–map similar to the neural network model used by Gerlee et. al. [42, 45] to study phenotypic plasticity. However, the CRN model would differ in that the determination of phenotypes could still be stochastic, permitting the study of environment–dependent bet–hedging strategies. The model of Gerlee et. al. [42, 45] is an extension of the classical concept of the reaction norm [34] to non– linear and higher dimensional functions (the output of which is then discretised to determine cell behaviour). The CRN model offers the natural next extension, breaking down the assumption of functionality within the reaction norm by introducing stochasticity, and brings the reaction norm concept more closely in line with biological reality. Thus, the theoretical extension of the genotype–phenotype mapping to a stochastic non–functional process can be further extended to account for environmental factors. Such an extension will bring the model more closely in line with the maxim of developmental biology, that both environment and genotype are equally important determining the phenotype ( “g + e = p”). Pigliucci [37] suggests, in his review of theoretical models of the genotype–phenotype mapping that we must build models which take both environment and genotype as equal partners in determining the phenotype and attempt to bridge the divide between developmental biology and the modern synthesis. The model introduced in this work represents a first step towards this goal but importantly offers something more than previous models that have set along this path — an attempt to account for the role of chance.

## Materials and Methods

### Stochastic Simulation of the CRNs

The chemical reaction networks are stochastically simulated using the Gillespie algorithm with tau–leaping [51, 52]. This algorithm determines a stochastic progression of the network and returns the sequence of reactions that occur and the times taken between them. As our model assumes that the molecular switch resolves sufficiently fast (in comparison to cell cycle times) that we may take it to be instantaneous, we ignore this timing information. As such, the abscissas of all figures showing stochastic simulations of chemical reaction networks presented in this work measure time discretely, in terms of the number of reactions that have occurred, instead of continuously. Probability distributions associating switching probability with initial conditions, such as those in row three of Figure 3 and Figure 4, are calculated by constructing the (absorbing) Markov chain on the state space of possible configurations.

### Population Dynamics

Consider an isogenic bet–hedging population with fixed GP–map and fixed genotype *g* corresponding to a probability *p* of giving rise to phenotype *A*. Note that as we wish to study evolutionary convergence to a one–phenotype strategy we may assume *p* ∈ (0, 1), as when *p* = 1 we will end our simulations. We use a discrete time difference equation model to simulate the population dynamics. The number of individuals of phenotype *A* and *B* in the population at discrete timestep t is denoted by x(*t*) = (*x*_*A*_(*t*), *x*_*B*_(*t*)). An individual of phenotype *A* (respectively *B*) survives and produces ƒ_*A*_ (resp. ƒ_*B*_) offspring on average over a single discrete timestep. Further, individuals of type *A* (resp. *B*) die with probability *d*_*A*_ (resp. *d*_*B*_) each time step. Note that we separate the death and birth parameters in order to more easily simulate the increase in death rate associated with a harsh environment (e.g. a drug treatment). The expected number of offspring individuals of either phenotype produce over a single timestep is given by w = (*w*_*A*_, *w*_*B*_), where we assume *w*_*A*_, *w*_*B*_ > 0. Each new offspring takes phenotype *A* with probability *p* (and *B* with probability 1 - *p*). We assume an unbounded population size. Thus, the projection matrix for the population dynamics is given by,

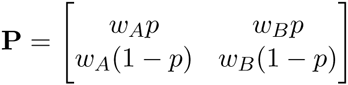
and the population distribution after one discrete time step is given by x(*t* + 1) = *Px*(*t*). As *p* ∈ (0, 1) the matrix P is positive and the Perron-Frobenius theorem, as applied to structured populations (see [65]), tells us that the dominant eigenvalue of the matrix *P* is in fact the net reproductive rate *r* for the hedging population. As we have only two phenotypes we can easily determine this dominant eigenvalue by solving,

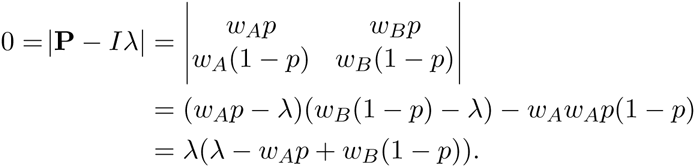

Hence the net reproductive rate, and equivalently the average fitness, of the population governed by *P* is given by *r* = *wAp* + *wB*(1 − *p*) and we know

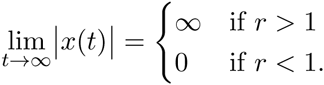

In the case *r* = 1, the population size remains constant and is dependent on the initial conditions of the system. This identity reveals the conditions in which a bet-hedging population will go extinct in terms of the probability *p* and the phenotype fitnesses w. The *fundamental theorem of demography* [66] tells us that if **v** is the *l*_1_-normalised left-eigenvector corresponding to eigenvalue *r* then,

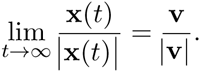

It is easy to verify then that x^*^ = (*p*, 1 − *p*) is the eigenvector satisfying this identity and is thus the equilibrium population distribution of phenotype *A* and phenotype *B*. Of course, this identity follows immediately from the definition of the bet-hedge - if every individual ever born has probability *p* of taking phenotype *A,* then the proportion with phenotype *A* must eventually (after any transient asymmetry introduced by the initial conditions has disappeared) be *p*. However, if we extend our model to allow the hedge to be dependent on the parent phenotype, to vary epigenetically or to be environmentally determined, then this identity will prove useful. It is worth noting that for a genotype *g* which gives a hedge *p*(*g*) the population dynamics are determined entirely by *p*(*g*) and are independent of the shape of the GP-mapping. It is only when we consider the difference in dynamics between populations of two different genotypes that the form of the GP-map begins to reveal itself.

### Invasion Dynamics

Our aim is to determine the long-term evolutionary trajectories of a population of cells endowed with different GP-maps. It is intractable to determine these trajectories through explicit simulation of the population alone. Instead we derive an analytic solution for the probability of a mutant genotype invading an existing isogenic population. Consider a large fixed-size population and assume that mutation is sufficiently rare (explicitly that the mutation rate *μ* and population size *N* satisfy *N*μ*logN* << 1) that we may consider strong-selection weak mutation (SSWM) evolutionary dynamics. Under these assumptions we can assume that the population is isogenic and that each time a new mutant appears in the population that mutant either fixes as the new population genotype or becomes extinct. We assume, as we did for our model of population dynamics, that reproduction occurs at discrete synchronized time steps and that each new individual first inherits the genotype *g,* with a small probability μ of mutation according to equation 2, and then the phenotype is determined according to a stochastic run of the CRN GP-map as discussed above.

Suppose a single mutant of genotype *g′* arises in an isogenic population of genotype *g* and denote by π(*g′*) the probability that this mutant reaches fixation at the population genotype. This probability is dependent on the phenotype of this initial mutant and is given by,

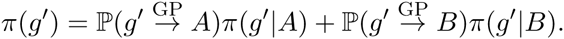

Denote π = (*p*(*g′*|*A*),*p*(*g′*|*B*)) and suppose that the population size, *N,* is sufficiently large that we may approximate it by the limit *N* → ∞. Assuming Wright-Fisher sampling for reproduction in our population, the value of π can be determined from the theory of branching processes. In particular, π can be calculated numerically as the solution to the equation 1 − π = *e*^−Mπ^ where, denoting the average fitness of the population by < *w* >,

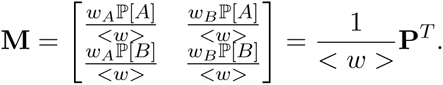

A proof of this identity, modified from the theory of viral quasispecies [67], is as follows. Under Wright-Fisher sampling the probability that a randomly chosen individual in the next generation is the offspring of a given individual of phenotype *i* in the current generation is 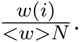. Thus, the probability that an individual is the offspring of a particular parent of phenotype *i* and genotype *g* and also has phenotype *r* is

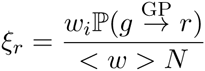
as not all offspring will have phenotype *r*. It follows that the probability that a given individual of phenotype *i* has precisely *k*_*r*_ offspring of phenotype *r* in the next generation is given by

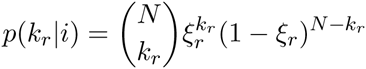

We can extend this argument to both phenotypes. The probability that an inidividual of phenotype *i* has *k*_*A*_ offspring of phenotype *A* and *k*_*B*_ offspring of phenotype *B* is

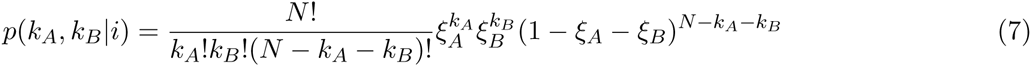

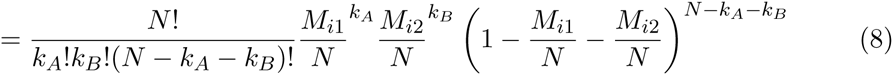
where the second equality follows from the definition of M. Assume now that the population size, *N,* is sufficiently large that we may approximate *p*(*k*_*A*_, *k*_*B*_|*i*) by taking the limit as *N*-→ ∞. This limit is a bivariate Poisson distribution

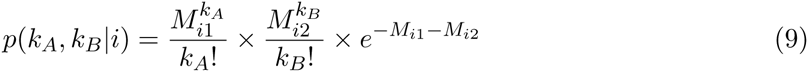

In the following derivation we will use the theory of branching processes [68]. This method, along with the assumption of infinite population size, restrict the resulting theory to fixation events of mutations which increase the average fitness of the population. Denote by *χ*_i_ the probability that the lineage of a single mutant individual with phenotype *i* in a population of average fitness < *w* > becomes extinct after finitely many generations. From the theory of branching processes we know that the vector of extinction probabilities ***χ*** = (*χ*_*A*_, *χ*_*B*_) satisfies ***χ*** = **f(*χ*)** where **f(z)** = (ƒ_*A*_(**z**), ƒ_*B*_(**z**)) is the probability generating function of the offspring probabilities *p*(*k*_*A*_, *k*_*B*_\*i*) given by,

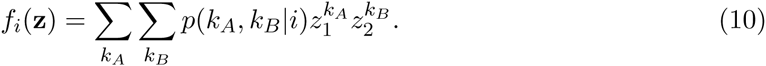

Substituting equation 9 into equation 10 we have

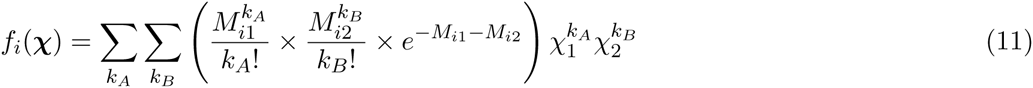

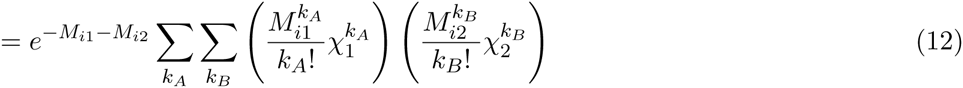

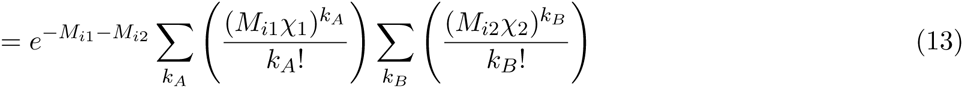

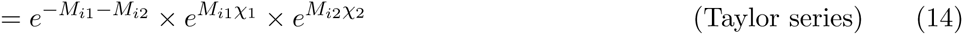

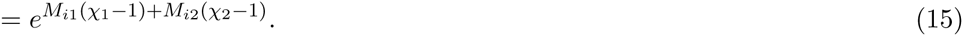

Taking the convention that *e*^x^ = (*e*^xi^,…, *e*^xn^) this gives

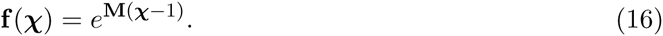

Now note that the probability of fixation of the mutant individual with phenotype *i* is precisely the probability that its lineage does not go extinct in finitely many generations, π = 1 − *χ*. Hence

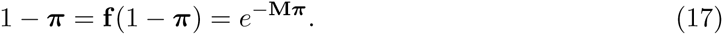

As the average fitness < *w >* can be determined by the population dynamics presented above, this equation can solved numerically. Figure 11 shows heatmaps of the likelihood that a single invader of with probability *p1* of phenotype *A* invades a resident population with probability *p2*. Heatmaps of invasion probability in terms of population genotypes are presented in Figure 7.

**Figure 11:**
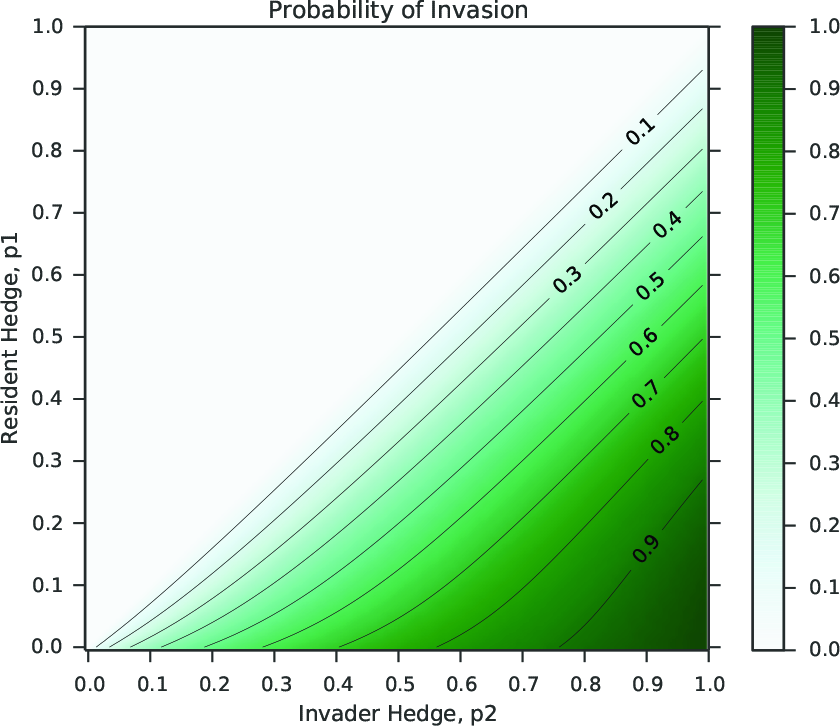
Invasion probabilities for a mutant with bet—hedging probability *p*_1_ into a resident population of phenotype *p*_2_. The parameters are those presented in Figure 3. Note that deleterious and neutral mutations cannot fix under our model of invasion dynamics, hence invasion in the case *p*_1_ ≤ *p*_2_ (above the antidiagonal of the plot) is impossible.

## Footnotes

1 This assumption can be justified by the observation that over these timescales either unmodelled mutations (such as mutations to the GP–map itself, to other genes governing the phenotypes A and B, or to other aspects of the phenotype) or unmodelled changes in the environment or ecosystem will occur, rendering our model unsuited to the situation.

